# Single-Cell Sequencing of Primate Preimplantation Embryos Reveals Chromosome Elimination Via Cellular Fragmentation and Blastomere Exclusion

**DOI:** 10.1101/241851

**Authors:** Brittany L. Daughtry, Jimi L. Rosenkrantz, Nathan H. Lazar, Suzanne S. Fei, Nash Redmayne, Kristof A. Torkenczy, Andrew Adey, Lina Gao, Byung Park, Kimberly A. Nevonen, Lucia Carbone, Shawn L. Chavez

**Author notes:** To whom correspondence should be addressed: Shawn L. Chavez, Ph.D., 505 NW 185th Avenue, Beaverton, OR 97006; Lucia Carbone, Ph.D., 3303 SW Bond Avenue, Portland, OR 97239.

## Abstract

Aneuploidy that arises during meiosis and/or mitosis is a major contributor to early embryo loss. We previously demonstrated that human preimplantation embryos encapsulate mis-segregated chromosomes into micronuclei while undergoing cellular fragmentation and that fragments can contain chromosomal material, but the source of this DNA was unknown. Here, we leveraged the use of a non-human primate model and single-cell DNA-sequencing (scDNA-seq) to examine the chromosomal content of 471 individual samples comprising 254 blastomeres, 42 polar bodies, and 175 cellular fragments from a large number (N=50) of disassembled rhesus cleavage-stage embryos. Our analysis revealed that the frequency of aneuploidy and micronucleation is conserved between humans and macaques and that cellular fragments encapsulate whole and/or partial chromosomes lost from blastomeres. Single-cell/fragment genotyping demonstrated that these chromosome-containing cellular fragments (CCFs) can be either maternal or paternal in origin and display DNA damage via double-stranded breaks. Chromosome breakage and abnormal cytokinesis resulted in reciprocal losses/gains at the terminal ends of chromosome arms, uniparental genome segregation, and mixoploidy between blastomeres. Combining time-lapse imaging with scDNA-seq, we also determined that multipolar divisions at the zygote or 2-cell stage generated chaotic aneuploidy encompassing a complex mixture of maternal and paternal chromosomes. Despite frequent chromosomal mis-segregation at the cleavage-stage, we show that CCFs and non-dividing aneuploid blastomeres exhibiting extensive DNA damage are prevented from incorporation at the blastocyst stage. These findings suggest that embryos respond to chromosomal errors by encapsulation into micronuclei, elimination by cellular fragmentation, and selection against highly aneuploid blastomeres to overcome chromosome instability during preimplantation development.

## INTRODUCTION

The demand for human *in vitro* fertilization (IVF) increases each year, but success rates as measured by live birth(s) have remained only ∼30-35% for decades (cdc.gov/art). One of the leading causes of IVF failure and embryo loss is the presence of unbalanced whole chromosome(s), or aneuploidy. Estimates of aneuploidy in IVF embryos via high-resolution techniques are 50-80%, including those from young, fertile couples and irrespective of embryonic stage (Vanneste et al. 2009a; Johnson et al. 2010; Chavez et al. 2012; Chow et al. 2014; Huang et al. 2014; Minasi et al. 2016). A similar efficiency (∼30-35%) is thought to arise from natural human pregnancies, with up to 70% of spontaneous miscarriages diagnosed as aneuploid (Miller et al. 1980; Wilcox et al. 1995; Zinaman et al. 1996; Ogasawara et al. 2000). Chromosomal mis-segregation in oocytes during meiosis has long been considered the primary reason for aneuploidy, especially in cases of advanced maternal age (Nagaoka et al. 2012). However, recent studies using comprehensive chromosome screening of all blastomeres in cleavage-stage human embryos established that mitotic errors occur at an equal or greater frequency and irrespective of maternal age (Vanneste et al. 2009a; Vanneste et al. 2009b; Johnson et al. 2010; Chavez et al. 2012; Chow et al. 2014; McCoy et al. 2015). Mitotic chromosome mis-segregation may not only lead to aneuploidy, but can also give rise to a mosaic embryo with different chromosomal copy number amongst cells. Euploid-aneuploid mosaic embryos may still result in the birth of healthy offspring upon transfer (Greco et al. 2015; Bolton et al. 2016; Fragouli et al. 2017), which suggests that corrective mechanisms exist to overcome chromosomal instability (CIN) during preimplantation development.

Another potential factor in the capacity of an IVF embryo to successfully implant is the timing and degree of cellular fragmentation, whereby cytoplasmic bodies pinch off of blastomeres during cytokinesis (Alikani et al. 1999; Antczak and Van Blerkom 1999). Distinct from cell death-induced DNA fragmentation (Hardy et al. 2001; Xu et al. 2001), cellular fragmentation also occurs naturally following *in vivo* human conceptions (Pereda and Croxatto 1978; Buster et al. 1985) and is not associated with maternal age (Wu et al. 2011). We previously demonstrated that cellular fragments can contain chromosomal material and that mis-segregated chromosomes are encapsulated into micronuclei during mitotic divisions (Chavez et al. 2012), but the parental source of this DNA and whether it originated from blastomeres was unknown. Chromosomes within somatic cell micronuclei display an increased propensity to undergo double-stranded breaks and structural rearrangements, which may be due to asynchrony in DNA replication timing between micronuclei and the primary nucleus (Crasta et al. 2012; Liu et al. 2018). A similar phenomenon has been proposed to occur in micronuclei of human embryos (Pellestor 2014; Pellestor et al. 2014), but a recent report suggests that mouse embryonic micronuclei do not rejoin the primary nucleus and instead undergo perpetual unilateral inheritance (Vazquez-Diez et al. 2016). Unlike humans, early cleavage-stage mouse embryos rarely exhibit aneuploidy, micronuclei, and cellular fragmentation even in sub-optimal culture conditions (Winston and Johnson 1992; Dozortsev et al. 1998; Lightfoot et al. 2006; Chavez et al. 2012; Macaulay et al. 2015; Bolton et al. 2016; Treff et al. 2016; Vazquez-Diez et al. 2016) and when micronuclei are induced experimentally mouse embryos undergo cell lysis rather than fragmentation (Chavez et al. 2014). At the late cleavage (morula) stage, however, ∼10% of mouse embryos have been shown to contain micronuclei and a similar number appeared between *in vivo* and IVF-derived embryos (Vazquez-Diez et al. 2016), to suggest that micronuclei formation is not a consequence of *in vitro* culture.

Previous studies with rhesus macaque embryos using DNA-fluorescent *in situ* hybridization (DNA-FISH) probes to human Chromosomes 13, 16, 18, X, and Y indicated that the incidence of aneuploidy in rhesus embryos is more comparable to human than mouse (Dupont et al. 2009a; Dupont et al. 2009b; Dupont et al. 2010). Given that only a few chromosomes were analyzed by low-resolution techniques, however, the actual percentage of rhesus embryos carrying chromosomal aberrations was unknown. Here, we used single-cell DNA-Sequencing (scDNA-Seq) to establish the frequency of whole and segmental chromosomal errors in 50 rhesus cleavage-stage embryos from the 2-cell to 14-cell stage. By reconstructing the chromosomal content of each cell and fragment, we investigated whether whole or partial chromosomes lost from blastomeres are sequestered into cellular fragments. We also examined the fate of cellular fragments beyond the cleavage-stage as well as embryo imaging behaviors or morphological features that might lead to their formation via time-lapse monitoring (TLM) of preimplantation development.

## RESULTS

### Incidence of micronucleation and cellular fragmentation is conserved between primates

To determine the aneuploidy and micronucleation frequency in rhesus cleavage-stage embryos, we developed an experimental approach utilizing scDNA-Seq and time-lapse imaging to non-invasively assess preimplantation development (**Fig. 1A**). Mature metaphase (MII) oocytes underwent conventional IVF and presumed zygotes with two polar bodies and/or pronuclei were analyzed by TLM to evaluate mitotic divisions, the absence or presence of cellular fragmentation (**Fig. 1B**), and other imaging behaviors and/or morphological features indicative of embryo chromosomal status. After ∼24-96 hours, cleavage-stage embryos (N=50) were disassembled into individual blastomeres, cellular fragments, and polar bodies if still present (**Fig. 1C**) for chromosomal copy number variation (CNV) analysis and single nucleotide polymorphism (SNP) genotyping. Another subset of intact embryos between the zygote and blastocyst stage were fixed and subjected to multi-color confocal imaging to assess micronuclei formation and DNA sequestration by cellular fragments. Lastly, an additional 92 rhesus embryos were allowed to proceed in development to evaluate the impact of micronuclei, fragmentation, and aneuploidy on embryonic arrest versus successful progression to the blastocyst stage (**Fig. 1D**).

**Figure 1.**
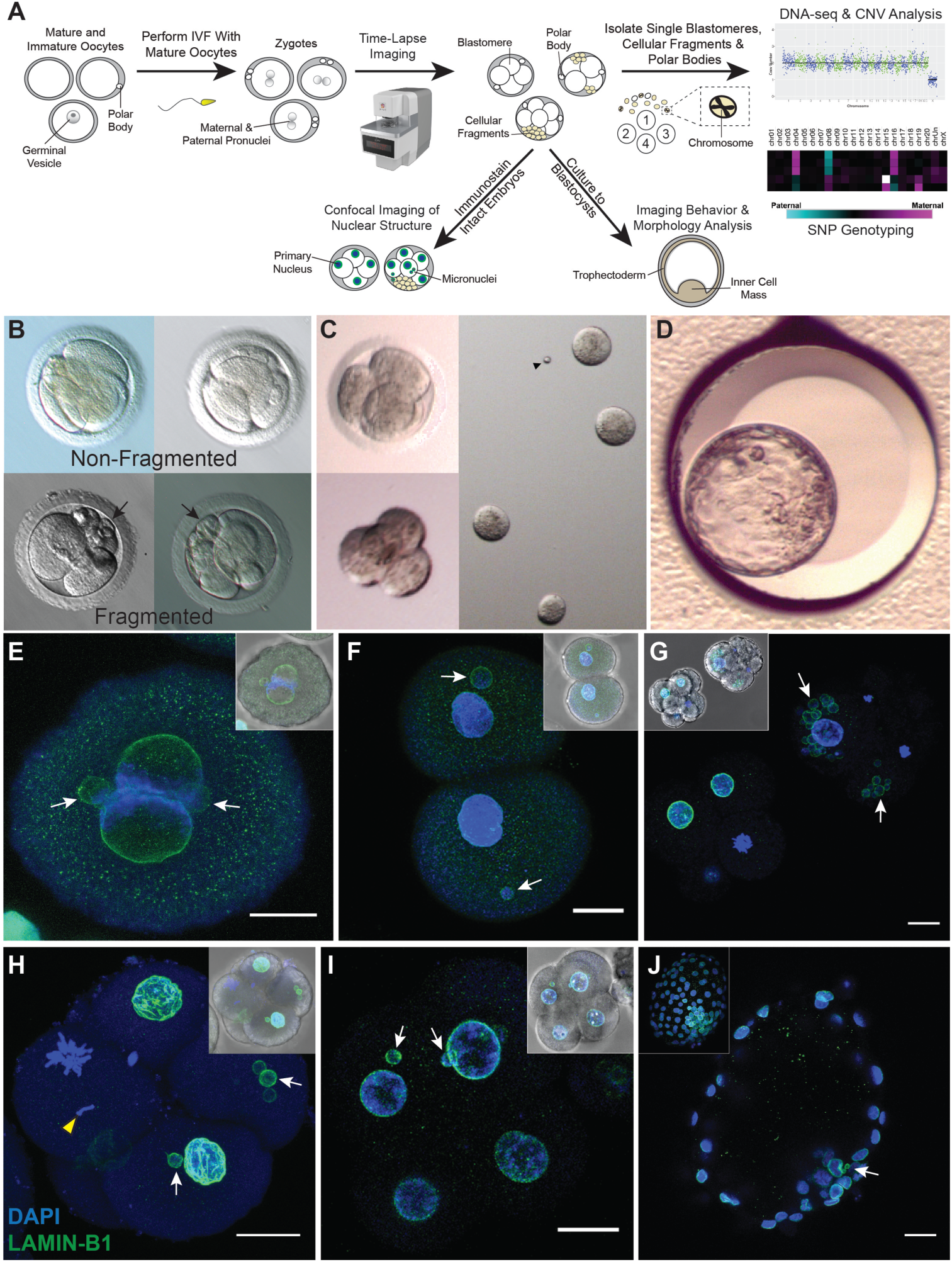
Approach for assessing CNV, micronuclei, and fragmentation dynamics in rhesus embryos. **(A)** Mature and immature oocytes were obtained from female rhesus macaques undergoing controlled ovarian stimulations. MII oocytes displaying one polar body were fertilized by conventional IVF with sperm from males. Early mitotic divisions and (**B**) the incidence of cellular fragmentation in presumptive zygotes (identified by two pronuclei and/or polar bodies) were analyzed by time-lapse imaging. **(C)** Cleavage-stage embryos were disassembled into individual blastomeres, cellular fragments, and/or polar bodies and analyzed by scDNA-Seq for CNV and SNP assessment. A subset of intact cleavage-stage embryos were fixed and immunostained for confocal imaging; **(D)** an additional group of embryos were cultured up to the blastocyst stage. **(E)** A zygote undergoing syngamy with two micronuclei (white arrows) using the nuclear envelope marker, LAMIN-B1 (green), and DAPI (blue) for nuclear DNA. **(F)** 2-cell embryo with one micronucleus in each blastomere. **(G)** Comparison of a fragmented (white arrowheads) cleavage-stage embryo with multiple micronuclei (right) and a non-fragmented 7-cell embryo (left). Single imaging plane of a Z-stacked **(H)** 5-cell embryo with a mis-segregated chromosome (yellow arrowhead) and micronuclei as well as a **(I)** 9-cell embryo exhibiting micronuclei in two blastomeres, but no visible cellular fragmentation. Insets show a brightfield image for reference. Scale bars: 25 μm. **(J)** Blastocyst with two micronuclei in the ICM; the inset shows the maximum intensity projection of the embryo.

Upon fixation and immunolabeling with the nuclear envelope marker, LAMIN-B1, we show that rhesus embryos contain micronuclei as early as the zygote (**Fig. 1E**) or 2-cell stage (**Fig. 1F**) and that the emergence of micronuclei is often concomitant with cellular fragmentation by the 4-cell stage (**Fig. 1G**). Some of these fragments encapsulate nuclear DNA positive for DAPI staining that is inconsistent with polar bodies, which contain condensed chromosomes and thought to degenerate within 24 hours of formation (Schmerler and Wessel 2011). While micronuclei did not appear until the 5-to 9-cell stage or later in certain embryos (**Fig. 1H**,**I**), we determined that the inner cell mass (ICM) of blastocysts might retain micronuclei (**Fig. 1J**). We also observed a condensed chromosome without nuclear envelope that was separate from the mitotic spindle (**Fig. 1H**), suggesting that mis-segregated chromosomes in embryonic micronuclei undergo condensation with nuclear envelope breakdown similar to chromosomes in primary nuclei. When all rhesus embryos were evaluated at high magnification prior to fixation or disassembly, we determined that >65% (N=129/196) of cleavage-stage embryos exhibit some degree of cellular fragmentation (**Supplemental Movie S1**). This indicates that the incidence of both micronucleation and cellular fragmentation is conserved between human and non-human primate embryos (Alikani et al. 1999; Antczak and Van Blerkom 1999).

### Rhesus cleavage-stage embryos are often aneuploid or mosaic due to mitotic errors

In order to perform scDNA-seq, embryos were disassembled into single cells and cellular fragments, the DNA in each sample was amplified, labeled with custom barcodes, PCR-validated using adapter sequences, and pooled for multiplex sequencing (**Supplemental Table S1**). To detect CNVs, we developed a bioinformatics pipeline that compares read counts in contiguous windows across the genome between embryonic samples and rhesus female euploid (42,XX) fibroblasts using a combination of Variable Non-Overlapping Windows and Circular Binary Segmentation (CBS) called VNOWC (**Supplemental Fig. S1**). We then employed a second custom bioinformatics pipeline that incorporated both CBS and Hidden Markov Model (HMM; Knouse et al. 2016) called CBS/HMM Intersect (CHI) to further validate CNV calls.

Utilizing the dual-pipeline bioinformatics strategy, we sequenced 471 individual samples from 50 whole rhesus cleavage-stage embryos up to the 14-cell stage (**Supplemental Table S2**), 49 of which contained DNA that successfully amplified (**Fig. 2A**). This included 254 blastomeres and 175 cellular fragments as well as a surprisingly large proportion of polar bodies (N=42/471) confirmed by SNP analysis as described below. Each blastomere or polar body was classified as euploid or aneuploid and the type of chromosomal error determined by the following criteria: (1) Meiotic errors were primarily identified by an aneuploid polar body and in the absence of polar bodies or presence of only one euploid polar body, it was considered meiotic if the same chromosome was affected in all sister blastomeres. (2) Mitotic errors were defined as different and/or reciprocal chromosome losses and gains between blastomeres with euploid polar bodies. (3) Chaotic aneuploidy was determined by greater than five randomly distributed chromosomes affected in one or more blastomeres. Based on the above criteria, 26.5% (N=13/49) of the embryos were comprised of only euploid blastomeres with no segmental errors, whereas 73.5% (N=36/49) contained at least one blastomere with whole and/or partial chromosome losses and gains (**Fig. 2A, Supplemental Table S2**). Further analysis revealed that 40.8% (N=20/49) of the embryos consisted of blastomeres that were all affected, while 26.5% (N=13/49) exhibited euploid-aneuploid mosaicism and 6.1% (N=3/49) were mosaic with segmental errors only. Both polar bodies were obtained from ∼20% (N=10/49) of the embryos and primarily at the early cleavage stages, but ∼74% (N=31/42) of those isolated were euploid. Thus, we were able to confidently call the inheritance of meiotic errors in 25% (N=9/36) and the occurrence of solely mitotic errors in 41.7% (N=15/36) of embryos, with the remaining 33.3% (N=12/36) either incurring both types of errors or unknown due to the complexity of chromosomal mosaicism (**Supplemental Table S3**). The incidence of chaotic aneuploidy was 28.6% (N=14/49) and this appeared to be mostly confined to embryos fertilized by a particular sperm donor (N=11/14). Representative examples of genome-wide chromosomal CNV plots from embryos with euploid, mosaic, aneuploid, and/or chaotic aneuploid blastomeres are shown in **Supplemental Figure S2**.

**Figure 2.**
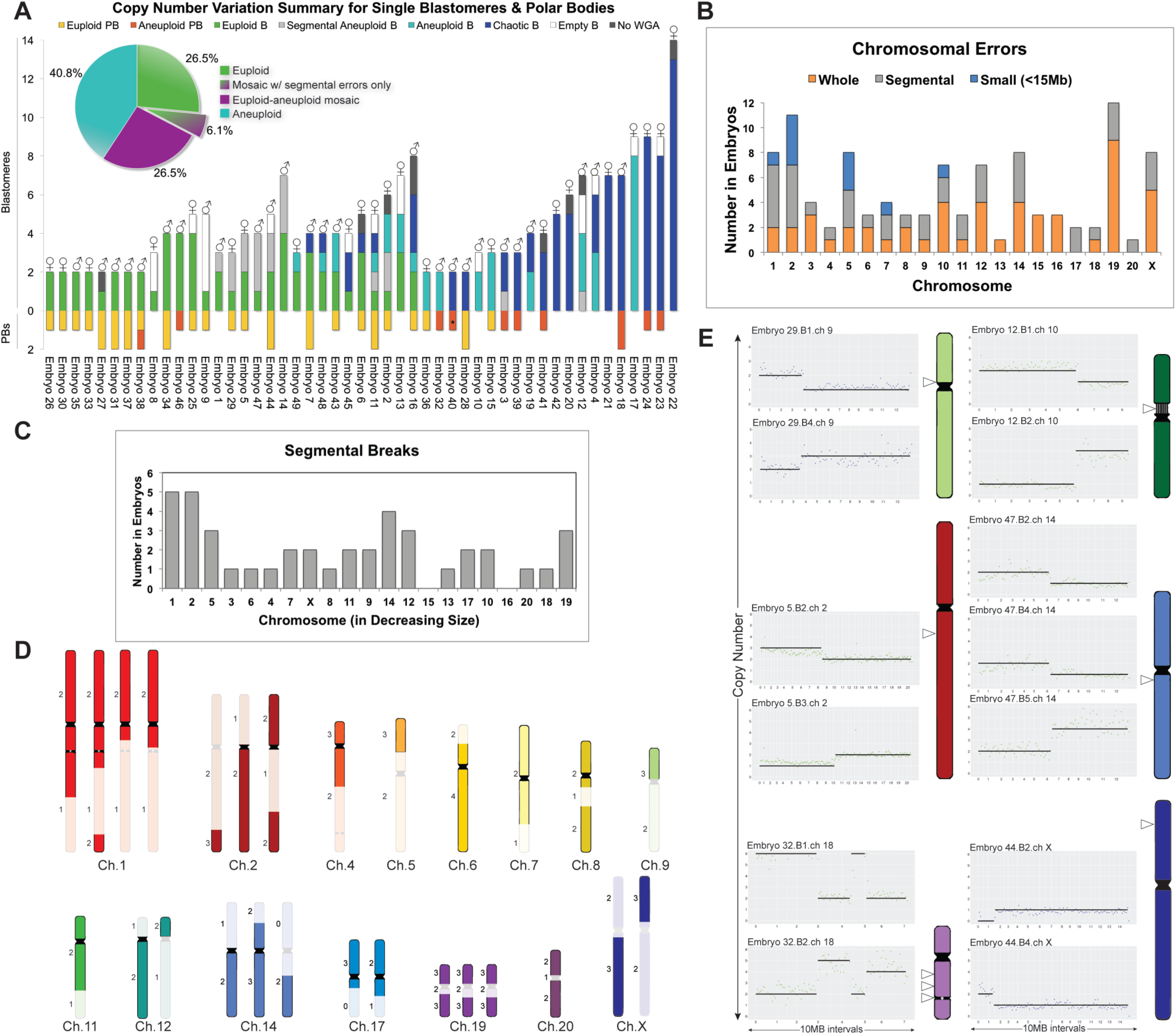
Assessment of whole and sub-chromosomal instability in blastomeres and polar bodies. **(A)** CNV summary of rhesus embryos (N=49) from the 2-to 14-cell stage analyzed by scDNA-Seq. Stacked bars represent euploid (yellow) and aneuploid (orange) polar bodies (PB); euploid (green), aneuploid (light blue), segmental aneuploid-only (purple), and chaotic aneuploid (dark blue) blastomeres (B); no WGA (gray); and empty blastomeres (white) detectable by high mitochondrial (mtDNA), but no genomic DNA reads. N=296 samples. Aneuploid PB containing segmental errors labeled with asterisk (*). ♂: Y Chromosome present ♀: X Chromosome(s) present. Percentage of euploid, aneuploid, or mosaic embryos with or without segmental errors only is shown in the pie chart (upper left corner). (**B**) Number of times chromosomes were affected by whole (orange) or segmental (gray) losses or gains. Small sub-chromosomal CNVs (<15Mb) shown in blue. **(C)** Graph showing that there was no significant association (p-value=0.1475) between the number of segmental breaks and chromosome size (Spearman’s correlation=0.3273). **(D)** Location of chromosomal breaks in embryos with segmental aneuploidy. Numbers to the left represent the blastomere copy number state. (**E**) CNV plots of six embryos, in which chromosomal breakage resulted in a reciprocal loss and gain of chromosome segments between blastomeres (left). Chromosome ideograms showing the approximate breakpoint locations (right; white arrowheads) of each embryo with reciprocal breaks. Vertical lines in Chromosome 10 delineate the nucleolus organizer region adjacent to the centromere (black) and the gray circle in Chromosome 18 designates the ancestral inactivated centromere.

### Reciprocal sub-chromosomal deletions and duplications indicate chromosome breakage

Excluding chaotic samples, we then assessed the frequency of whole, segmental, and small (<15Mb) sub-chromosomal errors by chromosome in the embryos (**Fig. 2B**). Chromosomes 1 and 2 were identified as the most highly susceptible to aneuploidy due to DNA breakage, which was not simply due to chromosome size (**Fig. 2C**), whereas Chromosome 19 experienced the greatest incidence of whole CIN. However, this chromosome is GC rich and when combined with scDNA-Seq as previously shown for human Chromosome 19, whole chromosomal losses and gains are difficult to distinguish from large segmental CNVs (Knouse et al. 2016). Chromosome 20 (corresponding to human Chromosome 16), on the other hand, was the least frequently affected by aneuploidy. Large segmental deletions, duplications, and amplifications were predominantly located at the terminal ends of chromosome arms (N=33/37; **Fig. 2D**). In a small proportion of embryos (16.7%; N=6/36), chromosomes underwent unbalanced rearrangements, in which the reciprocal chromosome segments were found in a sister blastomere (**Fig. 2E**). We determined that several of these breakpoints localized near existing centromeres or in the case of Chromosome 18, an inactivated ancient centromere (Ventura et al. 2007). The approximate location of breaks in Chromosomes 10 and 14 also aligned with corresponding fission or inversion evolutionary breakpoints, respectively, in the common primate ancestor (Ruiz-Herrera 2001).

### Cellular fragments may contain whole or partial chromosomes lost from blastomeres

Although reciprocal exchange of chromosomes between blastomeres was observed in some embryos, chromosome(s) were entirely lost from embryos in the majority of cases. Based on our previous findings of chromosomal material in cellular fragments of human embryos (Chavez et al. 2012), we hypothesized that the missing chromosome(s) had been sequestered during fragmentation. To test this, we sequenced 175 single fragments obtained from each fragmented rhesus embryo and found an instance in which both copies of Chromosome 9 and 12 lost from blastomeres were located in one of the cellular fragments (**Fig. 3A**). By similarly reconstructing the chromosomal content of each embryo via single-cell/-fragment DNA-seq, we observed additional examples of individual, multiple, and/or partial chromosomes in the fragments of other embryos (**Fig. 3B**). Maternal versus paternal SNP genotyping analysis, which is described in more detail below, revealed that chromosome-containing cellular fragments (CCFs) can originate from either the mother or father (**Fig. 3C**). There did not appear to be preferential sequestering of particular chromosomes, as both small and large chromosomes were affected and the partial chromosomes identified in fragments ranged in size from 6 to 85 Mb (**Fig. 3D**). Overall, we confirmed the presence of entire or portions of chromosomes in one or more CCFs in ∼18% (N=8/45) of fragmented embryos, but only ∼6.3% (N=11/175) of cellular fragments examined contained chromosomal material. In addition, 88% (N=7/8) of the embryos were aneuploid to varying degrees and it should be noted that only one of three blastomeres isolated from the single euploid embryo with CCFs successfully amplified for CNV analysis (**Fig. 3E**). Thus, mis-segregated chromosomes encapsulated within embryonic micronuclei not only persist or rejoin the primary nucleus, but may also be eliminated from the embryo upon cytoplasmic pinching of cellular fragments from blastomeres.

**Figure 3.**
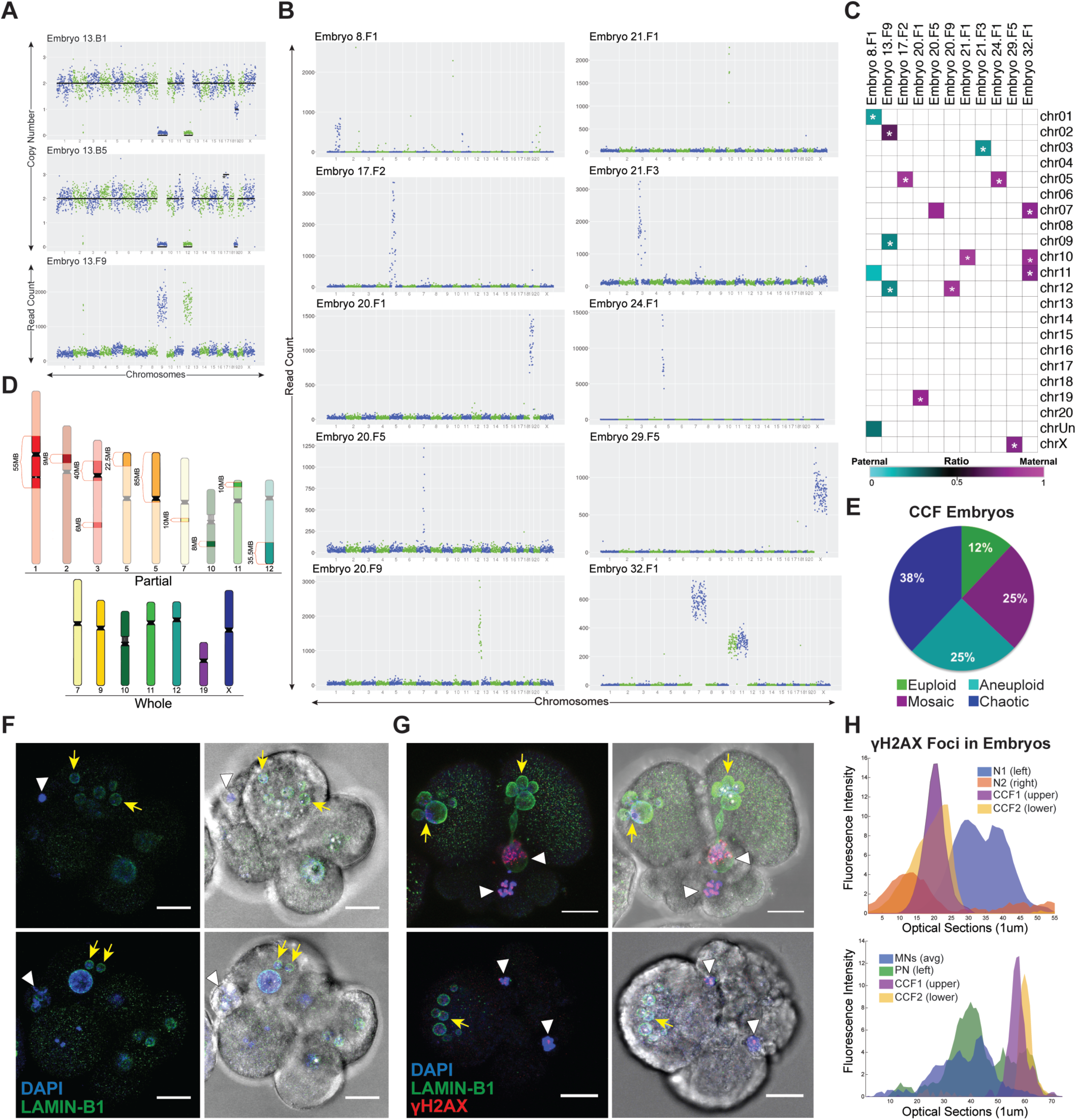
Micronuclei containing damaged chromosomes are eliminated via cellular fragmentation. **(A)** CNV plots demonstrating that chromosomes 9 and 12 lost from two blastomeres (top and middle; also missing 1-2 copies of chromosome 19 were detected in a cellular fragment (bottom) from the same embryo. **(B)** Additional examples of individual, multiple, and/or partial chromosomes in the fragments of rhesus embryos. **(C)** Heat map of maternal versus paternal SNP genotyping ratios showing that CCFs can originate from both the mother and father. White asterisk (*) demarcates significant p-values (p<9.1×10^−6^) for cumulative binomial test with Bonferroni correction. **(D)** Rhesus ideograms representing the whole (bottom) and partial chromosomes with approximate sizes highlighted that were detected in fragments. **(E)** Percent of embryos with CCFs (N=8 embryos) that were chaotic (blue), aneuploid (turquoise), mosaic (magenta), and euploid (green). **(F)** 8-cell and 7-cell (bottom) embryo with CCFs (white arrowheads) identified by DAPI (blue) signals that are similar in size to the LAMIN-B1 (green) positive micronuclei (yellow arrows) in adjacent blastomeres (left). Brightfield image (right) provided for reference. Scale bar: 25 μm. (**G)** 2-cell embryos with multiple cellular fragments and micronuclei immunostained for the double-stranded DNA break marker, γH2A.X (red), shows that the chromosomes within fragments are unstable and damaged. (**H**) γ-H2A.X fluorescence intensity measurements of primary nuclei (N or PN), micronuclei (MN), and CCFs from embryos to the immediate left.

### Chromosomes within cellular fragments are susceptible to DNA breaks and damage

Based on findings of whole chromosomes and/or chromosomal segments in cellular fragments from a minority of embryos and several reports of DNA fragility within the micronuclei of somatic cells (Crasta et al. 2012; Hatch et al. 2013; Liu et al. 2018), we next sought to determine whether the CCFs were susceptible to DNA damage and rapid degradation once separated from the primary nucleus. To accomplish this, we immunostained fragmented cleavage-stage embryos with LAMIN-B1 and gamma-H2A.X (γ-H2A.X), a marker of DNA damage and double-stranded breaks (Rogakou et al. 1998). Positive DAPI staining confirmed the presence of DNA in cellular fragments from rhesus embryos (**Fig. 3F**), but these CCFs appeared to have defective nuclear envelope as described for somatic cell micronuclei following DNA damage (Hatch et al. 2013; Liu et al. 2018). Multiple γ-H2A.X foci were detected in the micronuclei of blastomeres as well as in the DNA of cellular fragments (**Fig. 3G**) and by measuring γ-H2A.X fluorescence intensity, we determined that DNA damage was markedly increased in CCFs compared to both primary nuclei and micronuclei (**Fig. 3H**). Since DNA degradation and double-stranded DNA breaks are not distinguishable by γ-H2A.X immunosignals, however, it is also possible that sub-chromosomal segments were initially sequestered into cellular fragments rather than the product of DNA degradation.

### Parental contribution to aneuploidy is revealed by SNP genotyping analysis

It is generally accepted that polar bodies degenerate within 24 hours of extrusion from the oocyte or zygote (Schmerler and Wessel 2011), but we unexpectedly identified a large proportion of polar bodies in several embryos beyond the 2- to 4-cell stage (**Fig. 2A**). To validate their identity and distinguish them from cellular fragments, we isolated DNA from each of the parents whose gametes were used for IVF and performed whole-genome DNA-seq (**Supplemental Table S4**) for comparison of maternal versus paternal SNPs in all embryonic samples (**Supplemental Table S5**). The proportion of maternal SNPs was significantly different (p<1.88×10^−4^, binomial test) from the expected 50%, with an average of 80% of SNPs identified as maternal in origin, confirming polar body identity (**Fig. 4A**). SNP analysis was also used to assess the parental origins of all chromosomes in each blastomere. While the majority of euploid embryos were biparental (76.9%; N=10/13), three of these embryos contained blastomeres with chromosomes that were entirely from the mother (**Fig. 4B**) and either gynogenetic (embryos 8 and 25) or digynic triploid (embryo 9). In contrast, only ∼45% of the aneuploid embryos were biparental in origin (N=15/33) and the remaining embryos were gynogenetic (N=2/33), androgenetic (N=1/33), polyploid (N=11/33), or contained a mixture of uniparental, biparental, and triploid cells termed mixoploid (N=4/33; **Fig. 4C**, **Supplemental Fig. S3, Supplemental Table S5**). Further analysis revealed at least one case of a paternally contributed meiotic error (embryo 15; Chromosome 1 monosomy) and that the triploid embryos were comprised of two copies of maternal chromosomes and one copy of each paternal chromosome (**Fig. 4D**). When we compared the CCFs observed in **Figure 3E** to the embryos from which they arose via SNP genotyping, we determined that the blastomeres were biparental (N=1/8), gynogenetic (N=3/8), androgenetic (N=1/8), triploid (N=1/8), or mixoploid (N=2/8), suggesting that the production of CCFs is not associated with a certain type of chromosomal abnormality.

**Figure 4.**
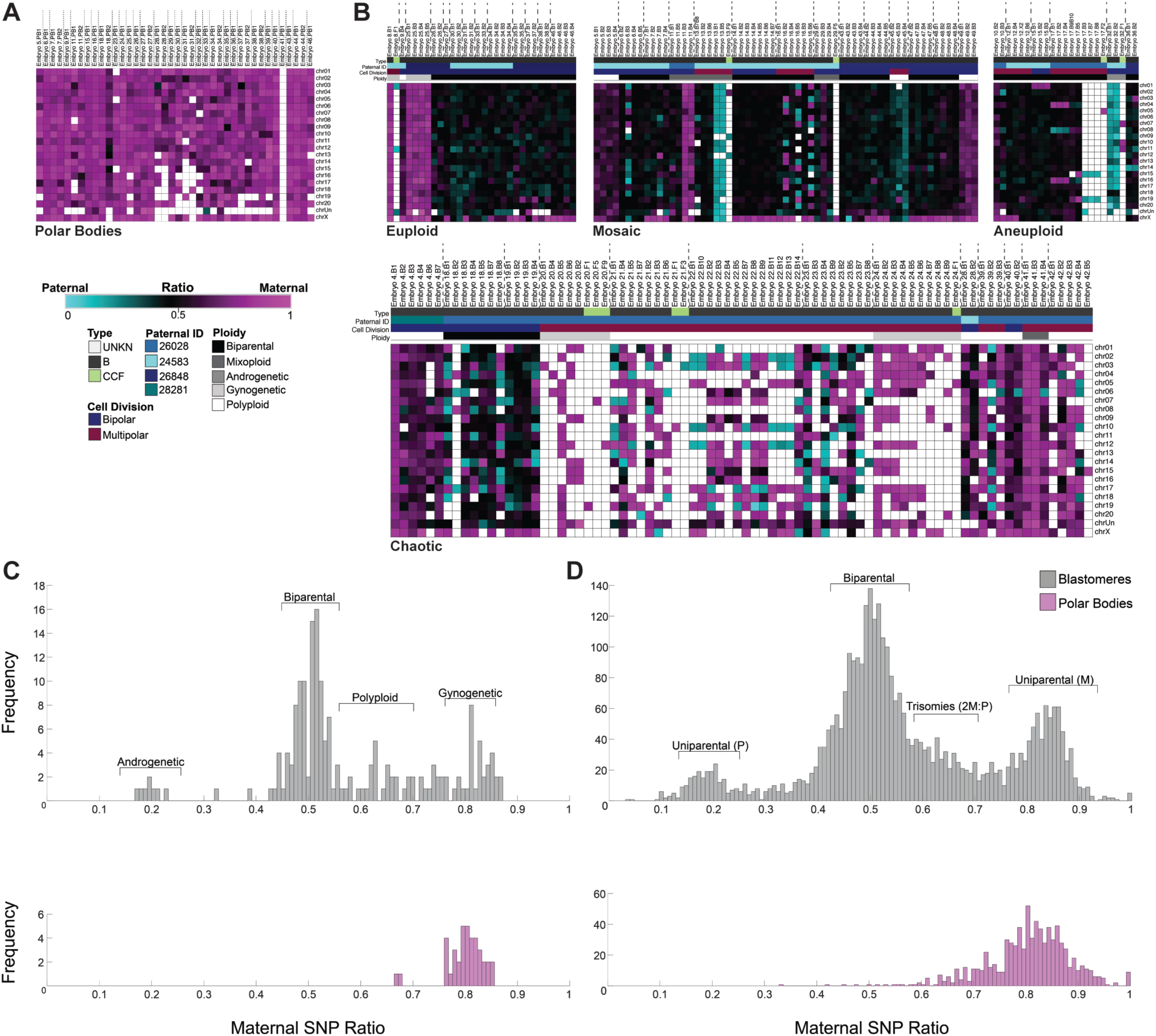
SNP profiling confirms polar bodies and reveals complexity in parental contribution to aneuploidy. **(A)** Heat map of SNP parentage ratios in presumptive polar bodies confirming their maternal origins and **(B)** of euploid, aneuploid, a mixture of euploid and aneuploid blastomeres (mosaic), or chaotic aneuploid embryos. Each embryo is separated by vertical dotted lines. Samples were further sorted based on the paternal donor, cell type, mitotic divisions, and overall embryo ploidy. Pink, blue, and black boxes indicate maternal, paternal, and biparental inheritance, respectively. White boxes show that either the chromosome was not detected or it could not be called with high confidence. **(C)** Histograms showing the distribution of SNP ratios across blastomeres (dark gray) and polar bodies (pink) in each sample revealed that the majority of rhesus embryos were biparental, but a small proportion were androgenetic, polyploid, or gynogenetic. **(D)** Frequency of SNPs ratios in histograms further stratified at an individual chromosome level.

### Multipolar divisions often lead to chromosome loss and chaotic aneuploidy

By combining scDNA-seq with TLM of embryos, we next sought to determine whether there were imaging behaviors indicative of chromosome loss from blastomeres and sequestration by cellular fragments. Indeed, the majority (N=6/8) of embryos with CCFs exhibited multipolar divisions at the 1-or 2-cell stage followed by cellular fragmentation (**Supplemental Movie S2**). When we evaluated higher order mitotic divisions in all embryos we determined that while only one of the 15 embryos with multipolar divisions was euploid, the remaining embryos were chromosomally abnormal and mainly chaotic aneuploid (N=8/15; **Fig. 5A**), with almost every blastomere affected (**Fig. 5B**). SNP analysis of parental ratios showed that all of the multipolar embryos with chaotic aneuploidy originated from the same sperm donor (**Supplemental Fig. S4**) and that multipolar divisions often resulted in a complex mixture of maternal and paternal chromosomes regardless of which male was used (**Fig. 5C**). In one of the multipolar embryos (**Fig. 5D**), we identified a loss of Chromosomes 4, 8, and 16 in three blastomeres and the reciprocal copies in two other blastomeres from the same embryo (**Fig. 5E,F**). These two blastomeres also contained only a single copy of Chromosome 19 and/or a complete loss of Chromosome 15, which were detected in additional cells that appeared unusual in shape and size upon disassembly. We were able to determine that this chromosomal complexity was due to a nondisjunction event that likely occurred during the multipolar division, resulting in paternal-only, maternal-only, or biparental contribution of certain chromosomes (**Fig. 5G**), all of which is depicted in **Figure 5H**.

**Figure 5.**
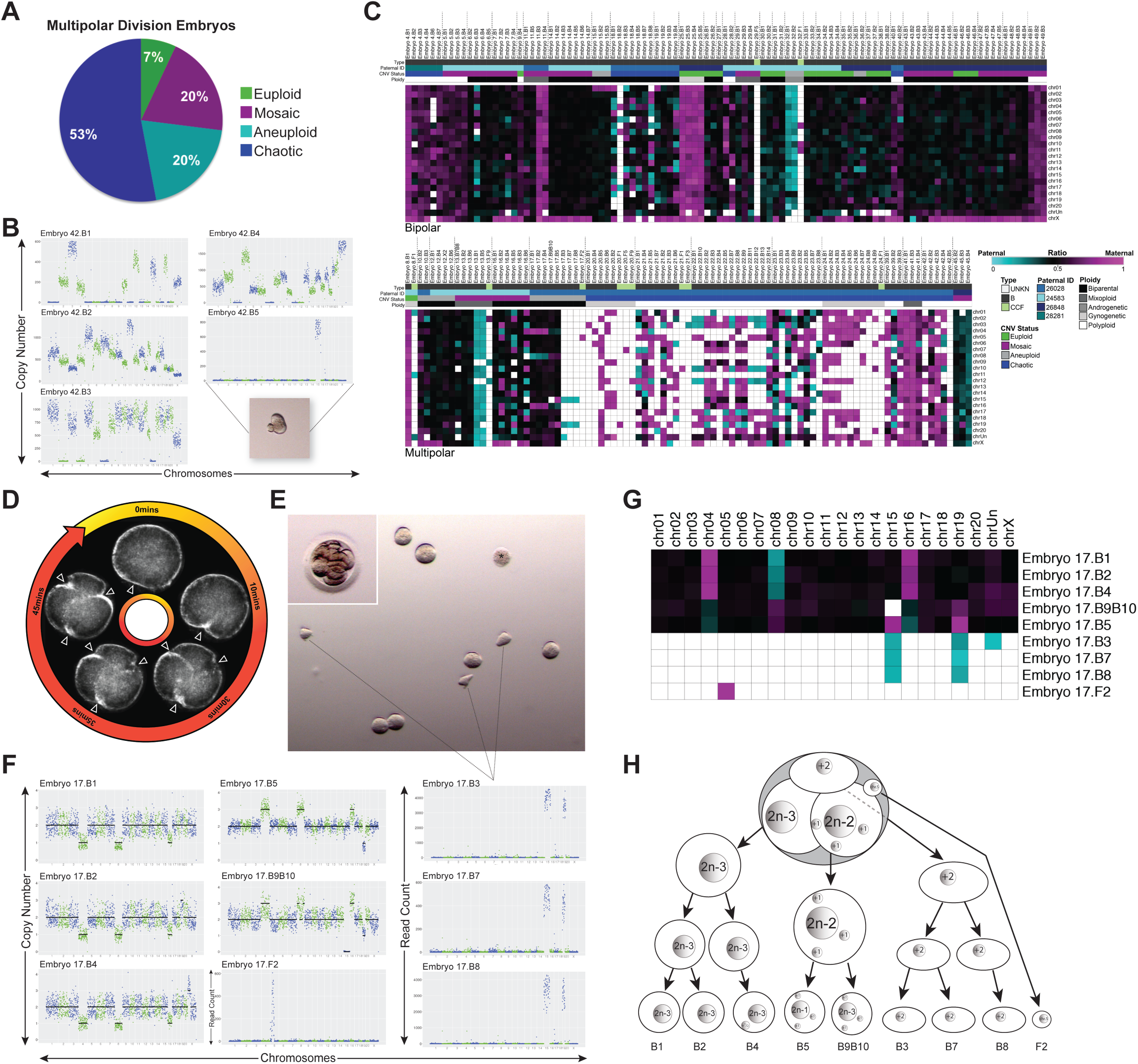
Multipolar divisions in embryos often result in chromosome loss and chaotic aneuploidy. **(A)** Ploidy status of rhesus embryos (N=15) with at least one multipolar division at the 1-or 2-cell stage. **(B)** CNV plots of blastomeres from an embryo, which underwent a multipolar 1^st^ division, showing chaotic aneuploidy in almost every cell. Inset is a stereomicroscope image of blastomere 5 containing only Chromosome 15 with a cellular fragment-like protrusion. **(C)** Heat map of SNP parentage ratios in embryos that underwent bipolar or multipolar cleavage(s) during the first three cell divisions. **(D)** Darkfield time-lapse images of a zygote undergoing a tripolar division. Arrowheads point to three simultaneous cleavage furrows. **(E)** Stereomicroscope image of the same embryo still intact (inset) and then disassembled. Blastomere 6 lysed and is demarcated with an asterisk (*). **(F)** The CNV plots of all whole blastomeres from the embryo showing multiple reciprocal chromosome losses and gains. Dotted lines indicate irregularly shaped blastomeres that each contained only two chromosomes. **(G)** Heat map of maternal versus paternal SNP ratios for this embryo delineates parental inheritance. **(H)** Schematic of the chromosome copy number state for each blastomere in this embryo based on the imaging and CNV analysis.

### Cellular fragments and non-dividing aneuploid blastomeres are excluded during blastocyst formation

In order to determine the impact of multipolar divisions and/or cellular fragmentation on subsequent rhesus preimplantation development, we monitored an additional 92 zygotes by TLM up to the blastocyst stage. While 42 of these embryos arrested prior to day 7 (**Supplemental Movie S3**; right), the remaining embryos formed blastocysts (**Supplemental Movie S3**; left), resulting in a typical blastocyst formation rate of ∼54% (N=50/92). Moreover, ∼18.5% (17/92) of the embryos underwent a multipolar division and ∼88% (N=15/17) of those arrested directly following the abnormal cytokinesis. The two multipolar embryos that still formed blastocysts exhibited a unique 1-to 4-cell symmetrical multipolar division without cellular fragmentation at the 1-or 2-cell stage. Of the blastomeres produced from these tetrapolar divisions, at least one cell ceased dividing and was confined to the blastocoel cavity upon blastocyst formation. We also observed the confinement of cellular fragments produced during the early cleavage stages to the perivitelline space of some blastocysts (**Fig. 6A**). Overall, we documented ten embryos with excluded blastomeres and/or cellular fragments that appeared at the 1-to 8-cell stage and persisted during the morula-to-blastocyst transition (**Fig. 6B**; **Supplemental Movie S4**). Numerous DAPI-positive nuclei were detected in the zona pellucida (ZP) of blastocysts that exhibited exclusion of cellular fragments after hatching (**Fig. 6C**). A large bi-nucleated cell with extensive DNA damage was also detected in one the blastocysts with excluded blastomere(s) via LAMIN-B1 and γH2A.X immunostaining (**Fig. 6D**). Since it was difficult to separate the excluded blastomeres from blastocysts once the blastocoel cavity formed, we disassembled other embryos with large non-dividing cells prior to or during morula compaction for scDNA-seq analysis. We determined that these excluded blastomeres were highly chaotic with multiple chromosomal losses and gains (**Fig. 6E**). SNP genotyping also showed that the chromosomes in excluded blastomeres were both maternal and paternal in origin (**Fig. 6F**). Because these blastomeres never divided again or in certain cases later lysed, this suggests that blastomere exclusion represents one mechanism by which an embryo can select against aneuploid cells during preimplantation development.

**Figure 6.**
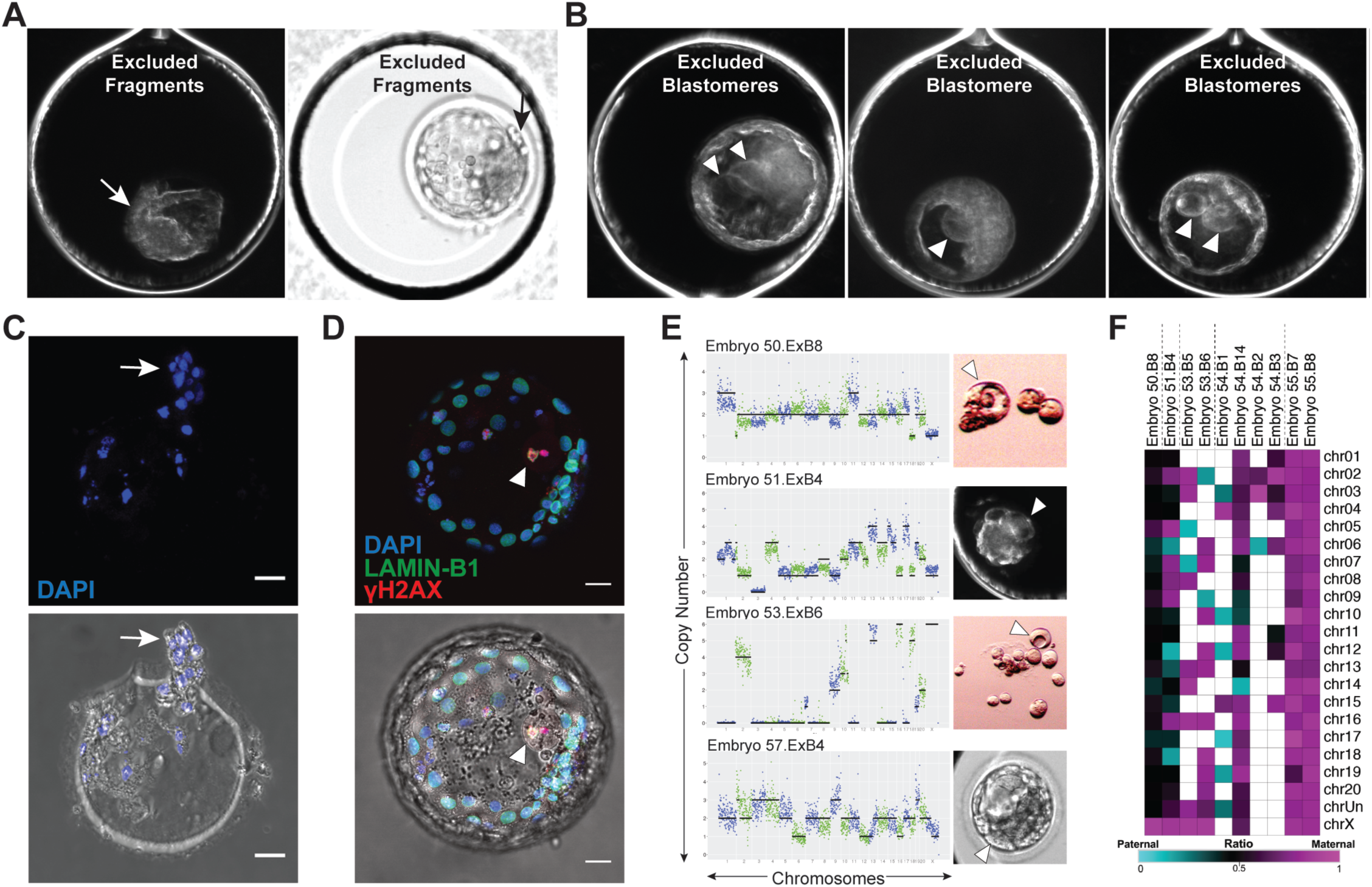
Cellular fragments and aneuploid blastomeres are excluded upon blastocyst formation. **(A)** Darkfield time-lapse image frames from four rhesus blastocysts exhibiting exclusion of several cellular fragments to the perivitelline space of the embryo (white arrow) or 1-2 non-dividing blastomeres to the blastocoel cavity (white arrowheads). **(B)** The zona pellucida of the blastocyst that exhibited cellular fragment exclusion with remaining DNA positive for DAPI (blue) staining following hatching (white arrow). **(C)** A blastocyst with blastomere exclusion immunostained for LAMIN-B1 (green) and γH2A.X (red) using DAPI as a marker for DNA. The large excluded blastomere appeared binucleated with strong γH2A.X signals (white arrowhead), indicating that double-stranded DNA breaks had occurred. Immunofluorescence with brightfield image overlay provided below. **(D)** Additional examples of large excluded blastomeres (white arrowheads; right) collected during the morula-to-blastocyst transition or at the blastocyst stage for sequencing. CNV analysis (left) determined that each excluded blastomere had chaotic aneuploidy. Scale bars: 25 μm. (**E**) Heat map of SNP maternal versus paternal ratios in excluded blastomeres shows varying parental origins.

## DISCUSSION

Established estimates of aneuploidy in human IVF embryos via whole-genome methods are 50-80% regardless of maternal age, fertility status, or embryonic stage and largely contribute to embryo arrest prior to the blastocyst stage (Vanneste et al. 2009a; Vanneste et al. 2009b; Johnson et al. 2010; Chavez et al. 2012; Chow et al. 2014; McCoy et al. 2015; Minasi et al. 2016). Here, we demonstrate that rhesus preimplantation embryos also have a high incidence of aneuploidy (73.5%) and chromosomal mosaicism due to meiotic and/or mitotic errors. Besides aneuploidy, we show that rhesus cleavage-stage embryos exhibit micronuclei formation, cellular fragmentation, and multipolar divisions at an equivalent frequency to human embryos (Alikani et al. 1999; Antczak and Van Blerkom 1999; Chavez et al. 2012; Hlinka et al. 2012; Chamayou et al. 2013; Ottolini et al. 2017). An examination of the time-mated breeding colony at our center during the same timeframe as this study (November-May; 2013-2017) revealed that of the confirmed ovulation and mating cases, 73.5% (N=200/272) did not result in a live birth to suggest a similar correlation between *in vitro* and *in vivo* conceptions. Based on all of the above, we argue that the rhesus monkey represents an ideal surrogate for studying the effects of human embryonic aneuploidy on normal preimplantation development (**Fig. 7A**).

**Figure 7.**
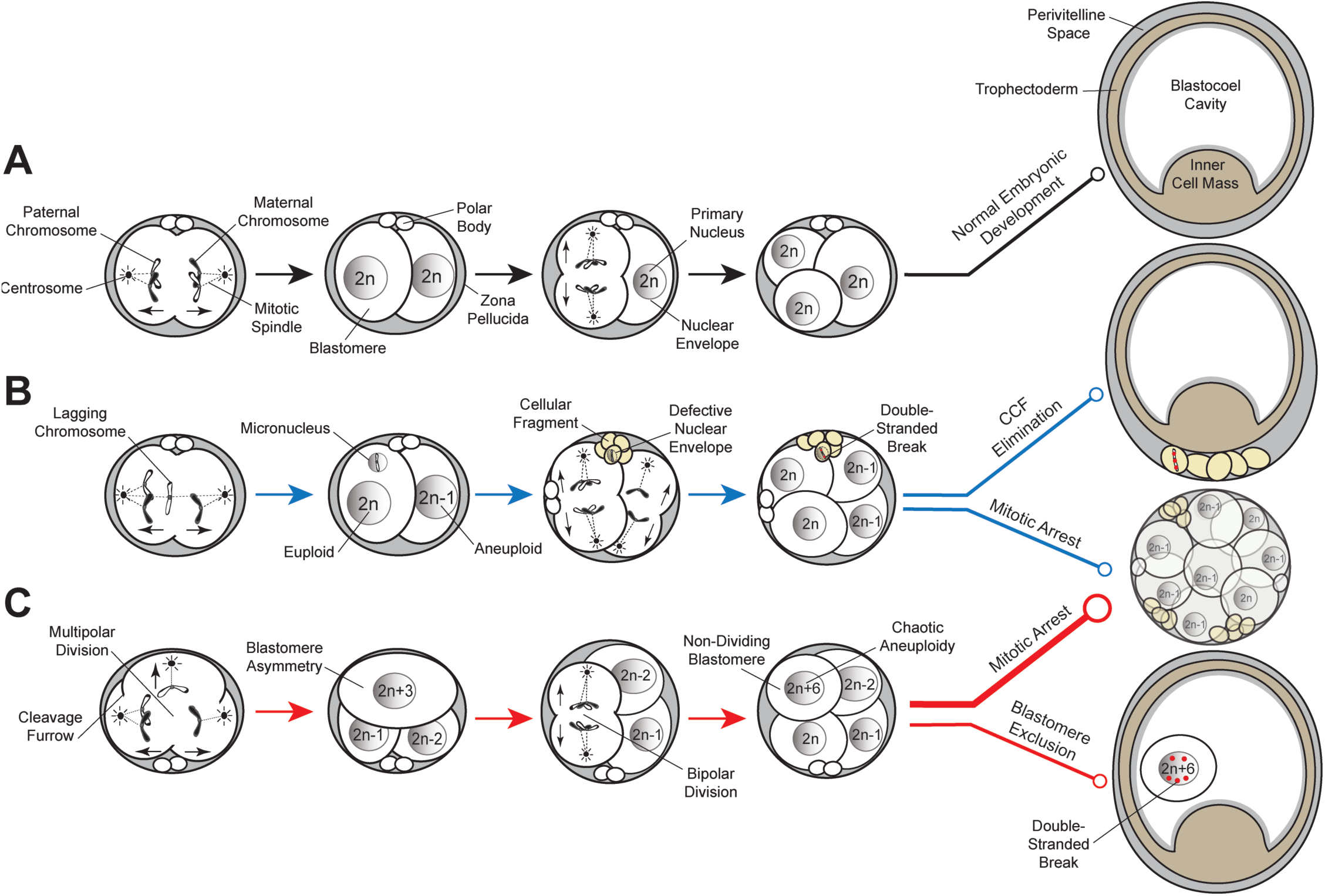
Proposed model of aneuploidy generation and potential resolution in embryos. **(A)** Simplified model of normal embryo development, whereby a euploid zygote undergoes proper chromosome segregation with bipolar cell divisions devoid of cellular fragmentation and blastomere exclusion (black lines). **(B)** A euploid zygote that contains a lagging chromosome from merotelic attachments during the first mitotic division becomes encapsulated in a micronucleus. Through the process of cellular fragmentation, the chromosome is eliminated from the blastomere, where it undergoes DNA damage in the form of double-stranded breaks due to defective nuclear envelope. This mosaic embryo is more likely to undergo mitotic arrest, but if it is able to progress in development to the blastocyst stage, the CCF may be sequestered to the perivitelline space (blue lines). **(C)** Multipolar cell divisions, including a tripolar cleavage occurring at the zygote stage, may also generate a mosaic embryo often with blastomere asymmetry. The vast majority of these embryos will be largely chromosomally abnormal with chaotic aneuploid blastomere(s) and eventually arrest at the ∼8-cell stage when a critical number of euploid blastomeres is not achieved (thick red line). Alternatively, mosaic embryos may still divide beyond embryonic genome activation and aneuploid blastomeres that fail to divide during this time will sustain DNA damage and become excluded to the blastocoel cavity upon blastocyst formation (thin red line).

To our knowledge, this is the first study to show that whole and/or partial chromosomes lost from blastomeres are encapsulated within cellular fragments and are highly unstable. Once separated from the primary nucleus, chromosomes within somatic cell micronuclei undergo DNA damage and double-stranded breaks due to defective nuclear envelope assembly (Crasta et al. 2012; Hatch et al. 2013; Liu et al. 2018). Because CCFs lacked nuclear envelope and we observed the greatest DNA damage in the sequestered chromosomes, this may explain why only a small number of cellular fragments contained intact DNA detectable by scDNA-seq. Chromothripsis, whereby chromosomes are “shattered” and rearranged in a single catastrophic event, also arises in somatic cell micronuclei following DNA damage (Zhang et al. 2015). The occurrence of chromothripsis in embryos has been suggested (Pellestor 2014; Pellestor et al. 2014), but not yet confirmed, due to the depth of genome coverage and large amplicon size required to accurately call structural variants by scDNA-seq (de Bourcy et al. 2014). We were limited by the same factors in this study, but did identify large segmental losses, duplications, and amplifications at the terminal ends of chromosome arms in rhesus blastomeres analogous to observations of terminal chromosome imbalances and rearrangements in human embryos (Vanneste et al. 2009a). Additional sequencing and bioinformatics approaches are required to delineate if there are structural differences in the chromosomes from embryonic micronuclei, cellular fragments, and excluded blastomeres, the latter of which may be more susceptible to chromothripsis. This assumption arises from the apparent requirement that the damaged chromosome(s) within somatic cell micronuclei be exposed to nucleoplasm of the primary nucleus before undergoing DNA repair and rearrangement (Zhang et al. 2015). Even if embryos do not undergo chromothripsis, we speculate that severely damaged DNA indicated by the extensive γH2A.X signals in CCFs and excluded blastomeres is selectively eliminated from the embryo to prevent further propagation of highly unstable chromosomes (**Fig. 7B**). Given the parallels between embryonic and somatic cell micronuclei as well as recent evidence that polyploid giant cancer cells may represent the somatic equivalent of blastomeres (Niu et al. 2017), additional embryo scDNA-seq studies may also inform the cancer field.

When we evaluated whether there were imaging behaviors indicative of chromosome sequestration by cellular fragments or blastomeres in embryos, there was a clear association between these two events and multipolar divisions. TLM has shown that ∼12% of human zygotes undergo multipolar divisions (Chamayou et al. 2013) and are less likely to form blastocysts and implant (Hlinka et al. 2012). Ottolini *et al*. also recently found that multipolar divisions occurring later in preimplantation development are highly correlated with human embryo arrest (Ottolini et al. 2017). In our study, almost all of the rhesus embryos with higher order divisions arrested prior to forming blastocysts and the two embryos that did progress underwent a 1-to 4-cell symmetrical cell division without fragmentation. These two embryos also exhibited blastomere exclusion during the morula-to-blastocyst transition to suggest that multipolar divisions might provide a mechanism to overcome aneuploidy under certain circumstances (**Fig. 7C**). This is supported by findings that some of the embryos with higher order divisions were either euploid with adjacent empty blastomeres or chromosomally mosaic. The most prevalent type of chromosomal abnormality observed in multipolar embryos was chaotic aneuploidy and all of these embryos shared a common sperm donor. Because the centrosome for the first mitotic division(s) is paternally inherited in most mammalian species except rodents (Sathananthan et al. 1991; Schatten et al. 1991), defective or supernumerary centrosomes from the sperm likely contributed to the higher order divisions. Moreover, sperm are also responsible for the activation of oocytes during fertilization (Whitaker 2006; Yoon et al. 2008), suggesting that premature oocyte activation might have also been a factor. Regardless of the underlying mechanism(s) and which male was used, multipolar divisions often generated a complex mixture of maternal and paternal chromosomes. Further investigation is required to determine if the abnormal cytokinesis resulting in the appearance of CCFs or excluded blastomeres exacerbates CIN or whether such events are a deliberate attempt to eliminate aberrant chromosomes and blastomeres from the embryo.

One of the most intriguing findings from the SNP analysis was the identification of a few euploid and a relatively large proportion of aneuploid embryos with cells that were derived from only one parent. This phenomenon, called uniparental genome segregation, has been described in bovine embryos at the zygote stage and was thought to be a consequence of *in vitro* oocyte maturation for fertilization (Destouni et al. 2016; Tsuiko et al. 2017). Our study is the first to show that uniparental genome segregation as well as mixoploidy also occurs in embryos following the maturation of oocytes *in vivo*, which is used in >98% of human IVF cycles (cdc.art/gov), and beyond the zygote stage. While it is fairly well established that gynogenetic and androgenetic embryos can result from IVF, we speculate that a similar percentage of human embryos with uniparental origins has not yet been reported given that current preimplantation genetic screening methods do not examine parental origins of aneuploidy unless SNP arrays are used. However, SNP arrays are rarely employed by clinics for CNV analysis due to high rates of allele drop-out and the need to include parental DNA to interpret SNPs (McCoy et al. 2015; McCoy et al. 2018). In summary, we show that chromosomal loss from primate preimplantation embryos is due to sequestration by cellular fragments and/or non-dividing blastomeres, which may denote mechanisms to surpass aneuploidy as embryos undergo implantation and continue in development. Additional work will be necessary to capture the formation and fate of micronuclei, CCFs, and excluded blastomeres in real-time at the single chromosome level and determine the molecular mechanisms underlying their production and resolution.

## METHODS

### Rhesus embryos

Oocytes were collected from adult female rhesus macaques of average maternal age (9.2 ± 2.3 years old) undergoing controlled ovarian stimulations (COS) as previously described (Stouffer and Zelinski-Wooten 2004). Briefly, 30 IU of recombinant human follicular stimulating hormone (FSH; donated from Organon, Roseland, NJ) was intramuscularly (IM) injected twice daily in cycling females at the start of menses for six consecutive days. On days 7 and 8, 30 IU of both FSH and recombinant human luteinizing hormone (LH; donated by Al Partlow) were co-injected twice each day. When estradiol levels reached greater than 200 pg/ml, females were IM injected with 0.1 ml/kg of Antide (donated by the Salk Institute, La Jolla, CA) the following day to block circulating gonadotropin-releasing hormone and prevent an endogenous LH surge. Approximately 36 hours prior to follicle aspiration, a single dose (1,100 IU) of human chorionic gonadotropin (hCG; EMD Serono Ovidrel®, Rockland, MA) was IM injected to initiate oocyte maturation.

Laparoscopic follicular aspirations were aseptically conducted on anesthetized animals using suction to obtain cumulus-oocyte complexes (COCs) collected in Tyrode’s albumin lactate pyruvate (TALP)-HEPES media with 0.3% bovine serum albumin (BSA; Sigma-Aldrich, St. Louis, MO) and 1% Heparin sodium salt solution. Oocytes were denuded by gentle micropipeting in TALP-HEPES media containing 0.3% BSA and 3% hyaluronidase (Sigma-Aldrich). Metaphase I (MI) and MII oocytes were incubated in TALP plus 0.3% BSA media at 37°C with 5% CO_2_ for 4-6 hours. Fresh semen was collected from 1 of 4 adult male rhesus monkeys of average paternal age (9.4 ± 1.5 years old) to minimize variability between sperm donors the same day as oocyte retrieval for conventional IVF. Mature MII oocytes were fertilized in 100 μL drops of TALP complete media (0.3% BSA and 0.006% sodium pyruvate) with 20 × 10^6^/ml “swim up” sperm further diluted 9:1 with activator solution [10.3mM caffeine (Sigma C-0750) and 10.2mM cAMP (Sigma D-0627) in saline] as previously described (Lanzendorf et al. 1990). Following IVF at 37°C with 5% CO_2_ for 14-16 hours, excess sperm were removed from the fertilized oocytes and visually assessed for two pronuclei and/or two polar bodies. The collection and preparation of oocytes and sperm was performed according to the approved Institutional Animal Care and Use Committee (IACUC) Assisted Reproductive Technologies (ART) Support Core protocol #0095 entitled, “Assisted Reproduction in Macaques.” All animal procedures adhered to the NIH Guide for the Care and Use of Laboratory Animals and were conducted under the direction and assistance of the veterinary staff and animal technicians in the Division of Comparative Medicine at ONPRC.

### Time-lapse imaging

Confirmed zygotes were transferred to custom Eeva^™^ 12-well polystyrene petri dishes (Progyny, Inc., New York, NY; formerly Auxogyn, Inc.) and cultured in 100 μL of one-step commercial media supplemented with 10% serum protein (LifeGlobal, Guildford, CT) under mineral oil (CopperSurgical, Trumbull, CT) at 37°C with 6% CO_2_, 5% O_2_ and 89% N_2_. Embryos were monitored with an Eeva™ darkfield 2.2.1 or bimodal (darkfield-brightfield) 2.3.5 time-lapse microscope system (Progyny, Inc) that fit into a small tri-gas incubator (Panasonic Healthcare, Japan) as previously described (Vera-Rodriguez et al. 2015). Images were taken every 5 min. with a 0.6 second (sec.) exposure time and each image was time stamped with a frame number and all images compiled into an AVI movie using FIJI software version 2.0.0 (NIH, Bethesda, MD). Morphological features such as cellular fragmentation, asymmetrical/multipolar division, and fragment/blastomere exclusion were examined and recorded. Media was changed every two days by transferring the embryos to a new imaging dish until collected for analysis.

### Embryo disassembly

The ZP was removed from each embryo by a ∼30 sec. exposure to warm Acidified Tyrode’s Solution (EMD Millipore, Temecula, CA) and washed with Ca^2+^ and Mg^2+^-free phosphate buffered saline (PBS). Cleavage-stage embryos were disaggregated into single cells, polar bodies, and cellular fragments if present with Quinn’s advantage Ca^2+^ and Mg^2+^-free medium with HEPES plus 10% human albumin (CooperSurgical) with or without brief exposure to warm 0.05% trypsin-EDTA (Thermo Fisher Scientific, Waltham, MA) as necessary. Each blastomere, polar body, and cellular fragment was washed three times with Ca^2+^ and Mg^2+^-free PBS to reduce carry over from cellular debris or residual DNA and collected individually in ∼2 μL of the PBS for transfer to a sterile Ultraflux™ PCR tube (VWR, Radnor, PA). All of the above was performed under a stereomicroscope equipped with a digital camera (Leica Microsystems, Buffalo Grove, IL), which has movie-making capabilities, to document the collection of every sample. Once tubed, samples were flash frozen on dry ice and stored at −80°C. Only embryos for which the disassembly process occurred effectively with no apparent loss of material were carried forward for library preparation and sequencing.

### Somatic cells

To develop the custom bioinformatics pipelines, human B-lymphocytes (GM12878, Coriell Institute, Camden, NJ) were used for CNV analysis as previously described (Vitak et al. 2017). Female human skin fibroblasts from patients with monosomy X or trisomy 21 (GM10179 and AG05024, respectively, Coriell Institute) as well as karyotypically normal female and male rhesus skin fibroblasts (AG08312 and AG08305, respectively, Coriell Institute) were obtained and grown in DMEM F12 medium (Gibco, Gaithersburg, MD) supplemented with 10% fetal bovine serum (FBS; Sigma-Aldrich). Cells were trypsinized with warm 0.05% trypsin-EDTA and the trypsin inactivated with DMEM F12 medium plus 10% FBS. The cell suspension was serially diluted in Ca^2+^/Mg^2+^-Free PBS until single cells were detected in drops for isolation in ∼2 μL of the PBS and transfer to the low-retention PCR tubes and storage at −80°C. Karyotyping of the human and rhesus primary fibroblasts (N=50 metaphase spreads per cell line) was performed by the OHSU Research Cytogenetics Laboratory. All cell lines showed low levels of karyotypic heterogeneity (**Supplemental Fig. S1A-D**) and a Chromosome 19 pericentric inversion was detected in the rhesus male fibroblasts (**Supplemental Fig. S1E**).

### DNA library preparation

Single blastomere, polar body, cellular fragment, and skin fibroblast samples underwent DNA extraction and whole genome amplification (WGA) using the PicoPLEX single-cell WGA Kit (Rubicon Genomics, Ann Arbor, MI) according to the manufacturer’s instructions with slight modifications. DNA was released from samples with cell extraction enzyme at 75°C for 10 min. and subsequently pre-amplified with PicoPLEX pre-amp enzyme and a primer mix via a 95°C hotstart for 2 min. and 12 cycles of gradient PCR. Pre-amplified DNA was further amplified with PicoPLEX amplification enzyme and 48 uniquely-indexed Illumina sequencing adapters provided by the kit or custom adapters with indices designed as previously described (Vitak et al. 2017). Adapter PCR amplification consisted of a 95°C hotstart for four min., four cycles of 95°C for 20 sec., 63°C for 25 sec., and 72°C for 40 sec. and seven cycles of 95°C for 20 sec. and 72°C for 55 sec. Libraries were quantified by Qubit High Sensitivity (HS) DNA assay (Life Technologies, Carlsbad, CA) and validated for sequencing by PCR amplification of the adaptor sequences using the PCR Primer Cocktail (Illumina, San Diego, CA) and visualized by 2% agarose gel electrophoresis. Only libraries with DNA quantities greater than the no-template controls were included in sequencing. 50 ng of DNA was prepared from each blastomere or fibroblast and 25 ng from both polar bodies and CCFs. Pooled libraries were purified with AMPure® XP beads (Beckman Coulter, Indianapolis, IN) and re-quantified by the Qubit HS DNA kit. Quality was assessed with a 2200 TapeStation and/or a 2100 Bioanalyzer (Agilent, Santa Clara, CA).

### Multiplex DNA sequencing

Pooled libraries of single skin fibroblasts, blastomeres, polar bodies, and cellular fragments were sequenced on Illumina platforms according to the following scheme: individual rhesus female (42XX) fibroblast controls (N=5) were first sequenced on an Illumina MiSeq using the 150bp paired-end protocol and generated a total of ∼9.66×10^6^ reads (∼1.93×10^6^ reads/sample). An additional 20 single fibroblasts were sequenced on an Illumina NextSeq 500 using a 75-cycle kit with a modified single-end workflow that incorporated 14 dark cycles at the start of the first read prior to the imaged cycles and generated a total of ∼34.5×10^6^ reads (∼1.72×10^6^ reads/sample). This step excluded the quasi-random priming sequences that are G-rich and lack a fluorophore for the two-color chemistry utilized by the NextSeq platform during cluster assignment. Based on the similar number of reads obtained between the two protocols, we then sequenced pools of 17 and 48 embryo samples on the Illumina MiSeq using the 150bp paired-end protocol, which generated a total of ∼30.7×10^6^ reads (∼1.81×10^6^ reads/sample), and ∼117×10^6^ reads (∼2.44×10^6^ reads/sample), respectively. Additional pools of 146 and 225 embryo samples were sequenced on the Illumina NextSeq again using our custom 75bp single-end protocol and designed indices and generated a total of ∼258×10^6^ reads (∼1.77×10^6^ reads/sample) and ∼227×10^6^ reads (∼1.01×10^6^ reads/cell line), respectively. A final pool of 81 embryo samples was sequenced on the Illumina NextSeq using the custom 75bp paired-end protocol and generated a total of ∼74.8×10^6^ reads (∼9.23×10^5^ reads/sample).

All raw sample reads were de-multiplexed and sequencing quality assessed with FastQC as previously described (Krueger et al. 2011). Illumina adapters were removed from raw reads with the sequence grooming tool, Cutadapt (Chen et al. 2014), which trimmed 15 bases on the 5’ end and five bases from the 3’ end, resulting in reads of 120 bp on average. Trimmed reads were aligned to the most recent rhesus genome reference, RheMac8 (Zimin et al. 2014), using the BWA-MEM option of the Burrows-Wheeler Alignment Tool (Salavert Torres et al. 2012) with default alignment parameters. To avoid read pile-ups due to common repeats, all repeat sequences were “masked” (converted to an “N”) in the RheMac8 reference genome using Repeat Masker (Tarailo-Graovac and Chen 2009). Resulting bam files were filtered to remove alignments with quality scores below 30 (Q<30) as well as alignment duplicates that were likely the result of PCR artifacts with the Samtools suite (Ramirez-Gonzalez et al. 2012). The average number of filtered and uniquely mapped sequencing reads in individual libraries was between 0.8 and 1.2 million (**Supplemental Table S1**).

### Parental DNA library construction and sequencing

Whole blood was obtained from the male and female rhesus macaque parents in K_2_EDTA vacutainer collection tubes (BD Diagnostics, Franklin Lakes, NJ) by the Colony Genetics Resource Core within the Primate Genetics Program at ONPRC. Parental DNA was extracted using the Gentra® Puregene® blood kit (Qiagen, Germantown, MD) according to the manufacturer’s protocol and stored at −80°C. Samples (1 ug) were fragmented using the Diagenode Bioruptor Pico (Denville, NJ) for a 300-400 base pair (bp) size selection. The NEBNext® DNA Library Prep Master Mix Set and Multiplex Oligos (NEB, Ipswich, MA) were then used to generate Illumina whole-genome sequencing libraries following the manufacturer’s protocol. Libraries were quantified with the Qubit HS DNA kit and size distribution assessed with the 2100 Bioanalyzer. Multiplexed libraries were sequenced at the Oregon State University Center for Genomic Research and Biocomputing on the HiSeq 3000 platform using the 150 paired-end protocol for a total of 2.84×10^9^ reads (1.56×10^8^ reads/sample). One parental sample (ID: 26129) was sequenced on the Illumina NextSeq using our custom 75bp paired-end protocol for a total of ∼3.50×10^8^ reads.

### Copy number variant calling

#### Variable Non-Overlapping Window CBS (VNOWC)

Although PCR-based WGA is more effective for CNV analysis than isothermal amplification (non-PCR)-based techniques as previously shown (de Bourcy et al. 2014), some read accumulation and dropout are expected from the PicoPLEX method. To model this in a single-cell euploid sample, five paired-end and twenty single-end individually sequenced rhesus 42XX fibroblast libraries were combined to form a paired-end and single-end reference, respectively. These reference samples were used to generate variable-sized windows with a constant number of expected reads per window. The same windowing method was applied to reads from forty single human fibroblast samples. We used the appropriate reference genome to calculate the observed-to-expected ratio of read counts in each window given the total number of mapped reads. Before applying this approach to embryos, the pipeline was trained and tested on rhesus male euploid (42,XY) cells, as well as human fibroblasts carrying known aneuploidies (trisomy 21 or monosomy X). R package DNAcopy (version 1.44.0) was used to perform circular binary segmentation (CBS) across each chromosome and identify putative copy number changes between windows (Olshen et al. 2004). Since the window ratios assume that reads are spread evenly across the genome, a recalibration step was necessary to correct for the expected number of reads given CNV. Samples with mostly empty windows were presumed to have a base copy number of one, while a base copy number of two was assumed for samples with reads in the majority of windows. Copy numbers provided by CBS were rounded to the nearest integer and plotted, along with the corrected ratios for each window. Our bioinformatics pipeline was able to successfully detect chromosome losses and gains in all rhesus and human fibroblast samples, including single cells (**Supplemental Fig. S1F**). Each call was confirmed with the use of Gingko (http://qb.cshl.edu/ginkgo), an open-source web tool for evaluating CNV in single-cells. To estimate false positive calls, reads from 10-cells of human trisomy 21 and monosomy X fibroblasts were subsampled 100 times to levels typical of a single-cell sample (1M, 500K, 250K, and 100K reads). These subsampled reads were mapped to four different window sizes (500, 1000, 2000, 4000 reads per window) for a total of 3,200 CNV runs. We calculated the unexpected calls for whole and segmental aneuploidies depending on the number of mapped reads and determined that window sizes containing 4,000 reads produced high confidence CNV calling (**Supplemental Fig. SG,H**).

#### CBS/HMM Intersect (CHI)

A bioinformatics pipeline for GC bias correction was also implemented as previously described (Vitak et al. 2017) that uses both the CBS and Hidden Markov Model (HMMcopy, version 3.3.0) (Ha et al. 2012) based on parameters determined previously (Knouse et al. 2016). All calls from the HMM and CBS methods generated CNV profiles of variable sized windows that were intersected on a window-by-window basis.

#### Integration of VNOWC and CHI Pipelines

Whole, segmental, and small CNV (<15 MB in length) calls generated by the VNOWC and CHI methods were compared to further validate CNV calling. The majority of CNV calls were shared between pipelines (N=150/177; 84.7%) and discordant calls detected between the VNOWC (N=18/177; 10.2%) and CHI (N=9/177; 5.1%) pipelines were primarily sub-chromosomal differences (**Supplemental Fig. S1I**). These discordant calls were further analyzed based on recently described criteria (Vitak et al. 2017). First, the discordant VNOWC plot was examined to determine if it represented the integer loss or gain for the majority of estimated copy number points within the appropriate range of windows. Each smoothed, GC corrected HMM CNV plot from the CHI method was then analyzed to determine if the majority of the Log2 transformed values were above 0.4 for a gain or below 0.35 for a loss. An analogous approach was used for CHI CNV plots to determine if the majority of the Log2 transformed values were above 1.32 for a gain or below 0.6 for a loss. If two or more out of these three criteria were satisfied, the CNV call was retained. Otherwise, we conservatively estimated that the remaining chromosomes (N=27) in question were euploid. Approximate DNA breakpoint locations were identified in rhesus chromosome ideograms adapted from Ventura et al. (Ventura et al. 2007), http://www.biologia.uniba.it/macaque/, http://www.biologia.uniba.it/primates/2-OWM/MMU/MMU, and by identifying the syntenic g-band interval in the UCSC human assembly, hg38.

### SNP parentage analysis

Whole genome reads from the parents and embryo samples were processed using a pipeline that followed the best practice recommendations of the Broad Institute’s Genome Analysis Toolkit (GATK; (McKenna et al. 2010; Van der Auwera et al. 2013), but adapted for rhesus macaque. Briefly, reads were trimmed using Trimmomatic (Bolger et al. 2014) and aligned to the Mmul_8.0.1 reference genome using BWA-MEM (Li and Durbin 2010). BAM post-processing included local re-alignment around indels using GATK and marking of duplicate reads using Picard tools (http://broadinstitute.github.io/picard). GATK’s HaplotypeCaller was used to produce gVCF files for each parent and embryo sample, followed by separate genotype calling using GenotypeGVCFs. SNPs identified in repetitive regions by Repeat Masker and with the exception of CCFs, those samples that had fewer than 10 SNPs per chromosome, were removed. The sequence data was processed and analyzed using DISCVR-Seq (https://github.com/bbimber/discvr-seq/wiki), a LabKey Server-based system (Nelson et al. 2011). To restrict the set of SNPs to only those of higher confidence, we required that there be at least two reads for each SNP. We further selected only the SNPs for which the two parents appeared to have opposite homozygous genotypes, meaning all reads matched one allele in one parent and the alternative allele in the other parent. While this reduced usable SNP numbers, combining multiple SNPs across each chromosome/fragment provided sufficient information to determine parentage. The ratio of maternal to paternal SNPs was used to assess parental inheritance and was visualized in heat maps by Morpheus (Broad Institute, Cambridge, MA) and in histograms (MATLAB R2016a).

### Immunofluorescence confocal imaging

ZP-free embryos were washed in PBS with 0.1% BSA and 0.1% Tween-20 (PBS-T; Calbiochem, San Diego, CA) and fixed with 4% paraformaldehyde in PBS (Alfa Aesar, Ward Hill, MA) for 20 min. at room temperature (RT). Once fixed, the embryos were washed with gentle shaking three times for a total of 15 min. in PBS-T to remove residual fixative. Embryos were permeabilized in 1% Triton-X (Calbiochem) for one hour at RT and washed in PBS-T as described above. To block non-specific binding, embryos were transferred to a 7% donkey serum (Jackson ImmunoResearch Laboratories, Inc., West Grove, PA)/PBS-T solution overnight at 4°C. LAMIN-B1 (ab16048, Abcam, Cambridge, MA) and γH2A.X (05-636, EMD Millipore) antibodies were diluted 1:1,000 and 1:100, respectively, in PBS-T with 1% donkey serum and embryos sequentially stained overnight at 4°C. Primary antibodies were detected using 488-or 647-conjugated donkey Alexa Fluor secondary antibodies (Thermo Fisher) to the appropriate species at a 1:250 dilution with 1% donkey serum in PBS-T at RT for 1 hour in the dark. Between antibody incubations, embryos were washed in PBS-T and the DNA stained with 1 μg/ml DAPI for 15 min. Immunofluorescence was visualized using glass bottom petri dishes (Mattek, Ashland, MA) and a Leica SP5 AOBS spectral confocal system. Z-stacks 1-5 μM apart were imaged one fluorophore at a time to avoid spectral overlap between channels. Stacked images and individual channels for each color were combined into composite images using FIJI (Schindelin et al. 2012). Mean fluorescence intensity of γ-H2A.X was also measured for each stack in FIJI by averaging and subtracting the mean fluorescent intensities of multiple cytoplasmic regions to filter out background noise and account for potential loss of signal in >50μm sections.

### Statistical methods

Significance of SNP parental ratios was examined using the cumulative binomial test with Bonferroni correction at p<0.05. Chromosome size versus number of segmental breaks was evaluated by Spearman’s correlation.

## DATA AND SOFTWARE ACCESS

All sequencing data has been deposited in the SRA under SRP121351. The VNOWC bioinformatics pipeline for calling chromosomal copy number is available at https://github.com/nathanlazar/Oocyte_CN. Ginkgo, the single-cell CNV pipeline used to validate VNOWC calls, can be found at http://qb.cshl.edu/ginkgo.

## ACKNOWLEDGEMENTS

We are grateful to the ONPRC ART Core for their assistance with oocyte and sperm collection, Colony Genetics Resource Core for providing the parental DNA, Molecular & Cellular Biology Core for the Mi-Seq runs, Biostatistics and Bioinformatics Core for the SNP analysis, and Imaging & Morphology Core for confocal microscopy (supported by Grant S10 RR024585), all under the auspices of the NIH/OD ONPRC core grant (P51 OD011092). The TLM part of the study would not be possible without the generosity and support from Auxogyn, Inc. We also thank the OHSU ExaCloud Cluster Computational Resource, which allowed us to perform the intensive large-scale data workflows. We give special thanks to Drs. C. Bishop, C. Hanna, and J. Hennebold as well as C. Ramsey for rhesus samples and/or embryology expertise, S. Vitak and A. Fields for sequencing support, as well as G. Schau and M. Yan for assistance in CNV pipeline development. We also thank members of the Chavez and Carbone labs for insightful discussions. B.L.D. was supported by the P.E.O. Scholar Award, N.L. Tartar Research Fellowship, and T32 Reproductive Biology NIH Training Grant (T32 HD007133). J.L.R. was supported by the Collins Medical Trust Foundation and Glenn/AFAR Scholarship for Research in the Biology of Aging. N.H.L. was supported by a fellowship from the National Library of Medicine Biomedical Informatics Training Grant (T15LM007088). This work was supported by the NIH/NICHD (R01HD086073-A1), National Centers for Translational Research in Reproduction and Infertility (NCTRI) pilot funds (P50 HD071836), Howard & Georgeanna Jones Foundation for Reproductive Medicine, Medical Research Foundation of Oregon, and Collins Medical Trust (to SLC).

## AUTHOR CONTRIBUTIONS

B.L.D. and S.L.C. designed the study, performed experiments, analyzed data, and wrote the manuscript. J.L.R. developed the single-cell library preparation and multiplex DNA-Seq approach. N.H.L. created and implemented the VNOWC CNV pipeline. S.S.F. aligned and genotyped the samples and generated SNP parentage summaries. N.R. performed TLM and provided support for the embryology and imaging experiments. K.A.T. developed and implemented the CHI CNV pipeline. A.A. provided single-cell sequencing expertise, the NextSeq runs, and human b-lymphocyte data. L.G. and B.P. provided biostatistical analysis for the SNP data and segmental breaks. K.N. prepared sequencing libraries from the parental DNA. L.C. assisted in the study design and interpretation of the sequencing results. All authors were involved in editing the manuscript.

